# A 20^th^-century *Cannabis* pollen increase in a Pyrenean varved record and its potential causes

**DOI:** 10.1101/2023.03.28.534506

**Authors:** Valentí Rull, Teresa Vegas-Vilarrúbia

## Abstract

The continuous, varved and absolutely dated sedimentary record of Lake Montcortès (Iberian Pyrenees) has provided evidence for a distinct and characteristic 20^th^ century (1980s) increase in *Cannabis* pollen (20C) that persists today. This event was coeval with the geographical shift of the hemp production center in the Iberian Peninsula from east to northeast (where Lake Montcortès lies), which was accompanied by a significant production increase. This increasing trend was fostered by the renewed interest of the paper industry in hemp and was promoted by the onset of European Union subsidies to hemp cultivation. Illegal cannabis crops could have also contributed to the *Cannabis* pollen increase, but sound evidence is still lacking. These preliminary conclusions should be reinforced by increasing the resolution of the current palynological record and modeling the dispersal of *Cannabis* pollen around the Montcortès region. More similar high-resolution records are needed to verify the geographical extent of the 20C event. Additionally, Lake Montcortès varved sediments are proposed as a suitable candidate to characterize the onset of the “Anthropocene” epoch (mid-20^th^ century), as currently defined by the Anthropocene Working Group.

## 1. Introduction

*Cannabis* is among the earliest human domesticates and has been an integral part of human life for millennia (Rull, 2022). Almost every part of this plant is able to provide a variety of products and services, such as textiles, cordage, fuel, paper, medicine, oil, food, drugs or biofuel, along with recreational, ritual, aesthetic and educational materials, among others (Clarke & Merlin, 2013, 2016; Gray et al., 2016). A database on the traditional uses of *Cannabis* (CANNUSE) is available at http://cannusedb.csic.es/ (last visited 24 March 2023) (Balant et al., 2021). Taxonomically, the genus *Cannabis* has a single species, *Cannabis sativa*, subdivided into two subspecies, *C. sativa* subsp. *sativa* and *C. sativa* subsp. *indica*. However, as usual in plants intensively used by humans, a number of new varieties have been created by artificial selection to adapt the plant to a diversity of biogeographical features and cultural needs (Barcaccia et al., 2020; Kovalchuk et al., 2020). Due to the illegal condition of this plant in most Western countries, the scientific study of *Cannabis* underwent a significant decline in the mid-20^th^ century (Duvall, 2014; Warf, 2014). However, the last decade has witnessed a revival of interest in *Cannabis*, especially in relation to the genetic and phytochemical characteristics of its different varieties, as well as its evolutionary origin, domestication and further diffusion. This resurgence is linked to a renewed interest in crop improvement, especially for medical purposes but also for other applications (Gray et al., 2016).

The Iberian Peninsula (IP) has been considered a keystone area for the natural and anthropogenic diffusion of *Cannabis* due to its peculiar biogeographical position and its diverse and dynamic cultural history (Rull, 2022). According to the recent meta-analysis of a database (CHIP, for *Cannabis-Humulus* Iberian Peninsula) with nearly 60 pollen records widespread across the IP (Rull et al., 2023), *Cannabis* would have entered the IP in its wild form by the Late Pleistocene (150-12 kyr BP). Domesticated forms would have arrived later in two main dispersal waves, the first in the Neolithic (7-5 kyr BP) and the second in the Middle Ages (500 CE onward), via continental (Europe) and maritime (Mediterranean) pathways. Maximum cultivation and hemp retting activities were recorded during the Modern Age (16^th^-19^th^ centuries). A unique *Cannabis* record was obtained in the annually-laminated (varved) sediments of Lake Montcortès, situated in the northeastern sector (Fig. 1), which contains the longest (last 3000 years), continuous (gap-free) and high-resolution (bidecadal) absolutely dated pollen sequence available for the Mediterranean region (Rull et al., 2021). In this record, *Cannabis* appeared in the early Middle Ages (∼600 CE) and has continued until today, showing significant variations, which have been correlated with historical sociocultural developments.

**Figure 1.**
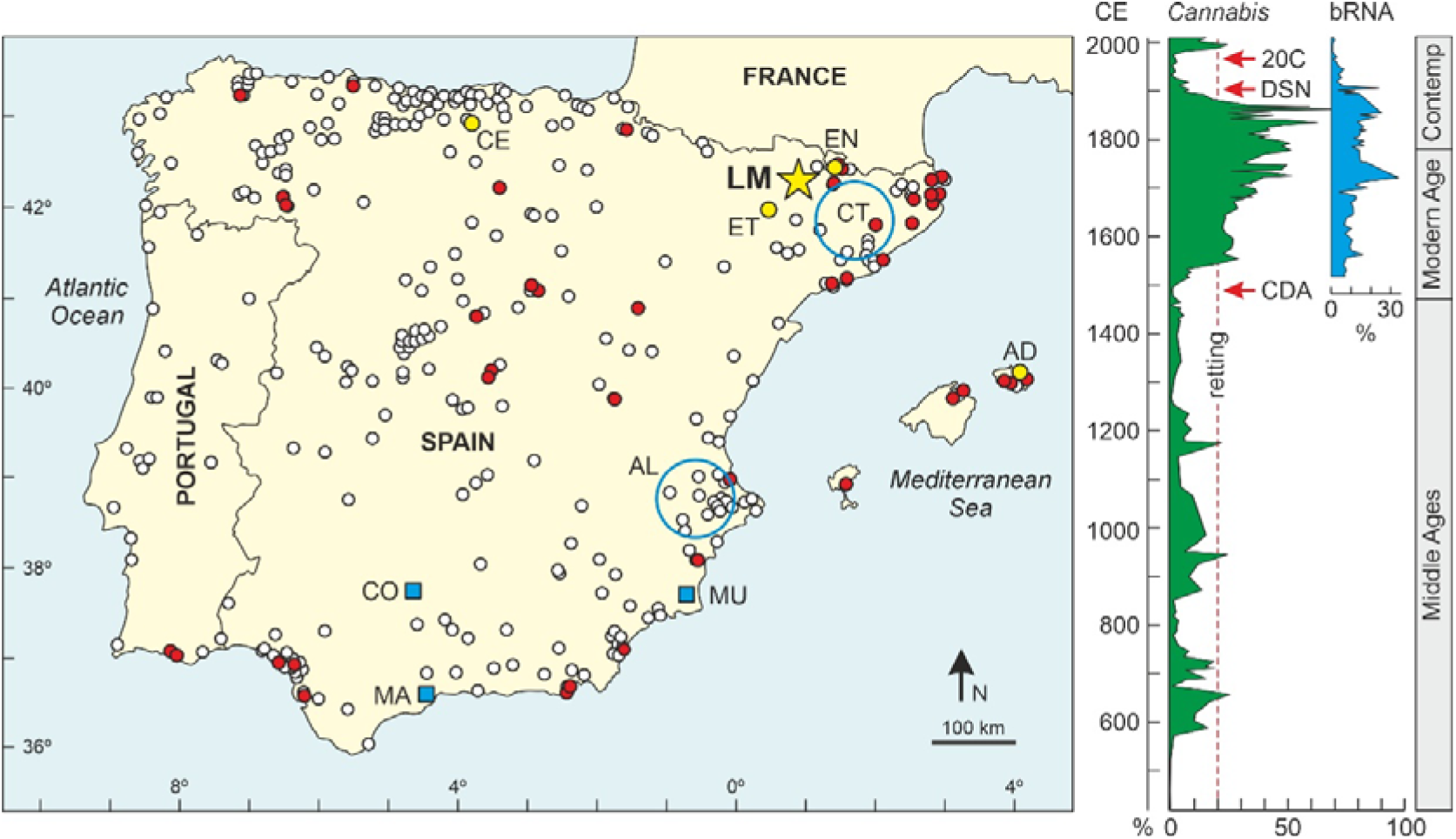
Left. Map of the Iberian Peninsula and the Balearic Islands (left), indicating the localities with published pollen records (white dots) (Carrión et al., 2022) and those containing *Cannabis* pollen (red dots) (Rull et al., 2023). Yellow dots indicate the sites with *Cannabis* records in the 20 ^th^century (AD, Alcúdia; CE, Cueto Espina; EN, Estanyons; ET, Estanya) (Rull et al., 2023). The location of Lake Montcortès (LM) is indicated by a yellow star. Blues boxes are the cities with aerobiological studies on *Cannabis* pollen mentioned in the text (CO, Córdoba; MA, Málaga; MU, Murcia). Blue circles are the main areas of hemp production, as explained in the text (AL, Alicante; CT, Catalonia). Right. *Cannabis* pollen curve (blue) and RNA from retting bacteria (bRNA) (blue) in percentage with respect to the total bacterial flora (Rull et al., 2021, 2022). CDA, Columbian discovery of America; DSN, dismantling of the Spanish navy; 20C, mid-20^th^ century *Cannabis* increase.

In the Middle Ages, *Cannabis* was below 20% of the pollen assemblages, which is considered the threshold for hemp retting (Rull et al., 2022). Therefore, the *Cannabis* pollen deposited in the lake would have been wind-transported from small-scale domestic hemp crops (Rull et al., 2021). The situation changed in the Modern Age, when hemp retting for fiber extraction seems to have been the main activity around the lake. By those times, hemp fiber extraction was accomplished by immersion of plants in lakes and ponds, where autochthonous pectinolytic bacteria naturally degraded the cementing compounds to separate the fiber from the stalk (Akin, 2013). The immersion of male hemp plants in lakes caused the pollen to be released and incorporated into sediments in larger amounts than expected if the only source of this pollen was wind transport from local and/or regional crops. The dramatic Modern-Age hemp pollen increase in Montcortès has been linked to the general increase in the hemp industry to meet the needs (mainly sails and ropes) of the Spanish navy, between the Columbian discovery of America (1492 CE) and the decadence of the Spanish empire, with the ensuing dismantling of its royal navy (1834 CE) (Rull & Vegas-Vilarrúbia, 2014). In Lake Montcortès, hemp-retting activity during this ∼350-y period was demonstrated not only by *Cannabis* pollen values over 20% (Fig. 1) but also by the identification of sedimentary RNA from bacteria actually responsible for the retting process (Rull et al., 2022).

During the first half of the 20^th^ century, hemp retting ceased in Lake Montcortès (Rull et al., 2022), and *Cannabis* pollen dropped to minimum historical values, coinciding with a major depopulation (60% reduction) of the Pallars region, where the lake lies, caused by the crisis of the subsistence economy and the massive emigration to industrial cities (Farràs, 2005). The *Cannabis* pollen suddenly increased again to values close to the retting threshold during the second half of the 20^th^ century (Fig. 1). However, bacterial RNA (bRNA) analyses show that local hemp retting never returned (Rull et al., 2022), and therefore, the source for this pollen rise – called here the 20^th^ century *Cannabis* increase (20C) – should be sought in other activities. Although some potential causes for the 20C event have been suggested (Trapote et al., 2018; Rull et al., 2022), the issue has not yet been addressed in depth. This paper analyzes the trends of the last century and the patterns of pollen deposition in recent years to discuss the potential causes of the 20C in relation to the main developments of the hemp industry in the IP.

## 2. Material and methods

The raw data used in this study are from Rull et al. (2017, 2021, 2022) and Trapote et al. (2018) and are available at https://data.mendeley.com/datasets/mr4h3×7×35 (pollen) and the Supplementary Material of Rull et al. (2022) (bRNA). In this analysis, *Cannabis* has been excluded from the pollen sum because the parent plant is not part of the local/regional vegetation (Carreras et al., 2005-2006; Mercadé et al., 2013). Pollen percentages have been recalculated and the diagrams have ben redrawn, with a focus on the 20^th^ century (1896-2013) and a 2-year modern-analog study (2014-2015) using sediment traps submerged in the lake. Details on the study site and the methods used are available in former papers. The age-depth model was based on varve counting (Corella et al., 2014, 2016), the palynological methods are described in detail in Trapote et al. (2018) and Rull et al. (2017, 2021), and the methods for bRNA analysis are available at Rull et al. (2022).

## 3. Results

The pollen trends of roughly the last century (1896-2013) are shown in Fig. 2, where it can be seen that the regional vegetation, consisting mainly of a mosaic of *Pinus* and *Quercus* forests intermingled with grasslands and crops, has experienced little variation. The main vegetation shift occurred in the second part of the 17^th^ century, when a conspicuous expansion of conifer forests coincided with a significant decline in olive groves, paralleled by a meaningful increase in total plant cover, as manifested in pollen accumulation rates (PAR). Regarding *Cannabis*, the main shifts occurred in the 1920s and 1980s. The first was a decline that signified the end of the hemp retting decline initiated in the late 19^th^ century, as manifested in the bRNA curve. This *Cannabis* decline is only evident in percentage units, whereas influx values show that this pollen was virtually absent from the beginning of the century. The second increase is the 20C event, when *Cannabis* increased and attained values over 30% in the 1990s, which is beyond the threshold for hemp retting (Rull et al., 2022). Influx values confirm that this increment was actually an increase in the amount of *Cannabis* pollen reaching the lake, rather than a percentage artifact. The 20C persists today, as indicated by a study of modern pollen sedimentation using sediment traps submerged in the lake (Rull et al., 2017). Indeed, the average amount of *Cannabis* pollen settled during 2014 and 2015 was slightly above 11%, with maxima over the retting threshold in winter and fall (Fig. 3). However, retting practices are no longer developed in the lake.

**Figure 2.**
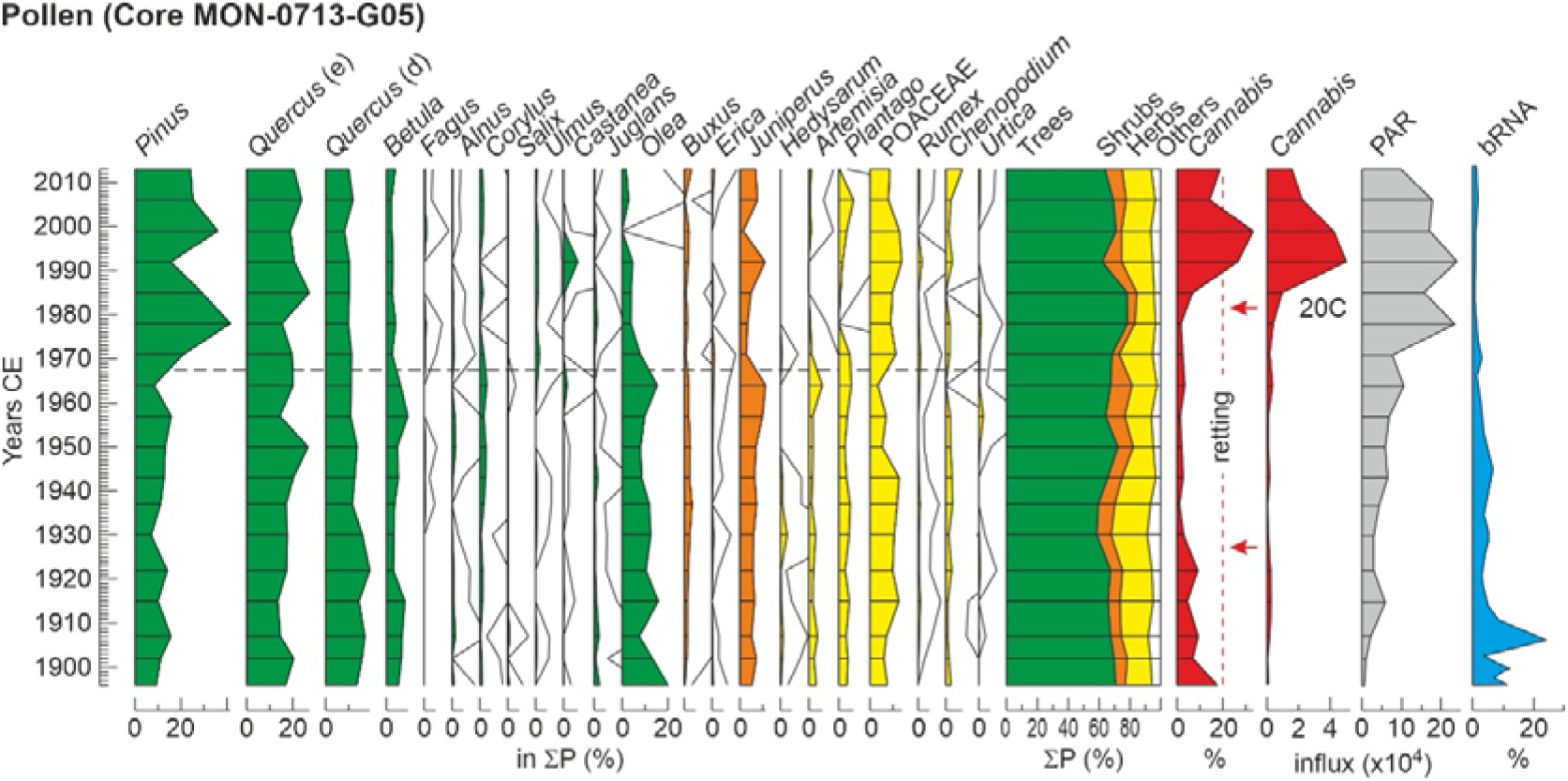
Percentage pollen diagram corresponding to the 20 ^th^century at subdecadal (∼5-year) resolution. Influx values (grains/particles cm^-2^-y ^-1^) for *Cannabis* and pollen accumulation rates (PAR) are also provided. RNA from retting bacteria (bRNA) is represented in percentage with respect to the total bacterial flora (Rull et al., 2022). Solid lines are x10 exaggeration. Red arrows represent the major shifts in *Cannabis* pollen percentage. ∑P, pollen sum.

**Figure 3.**
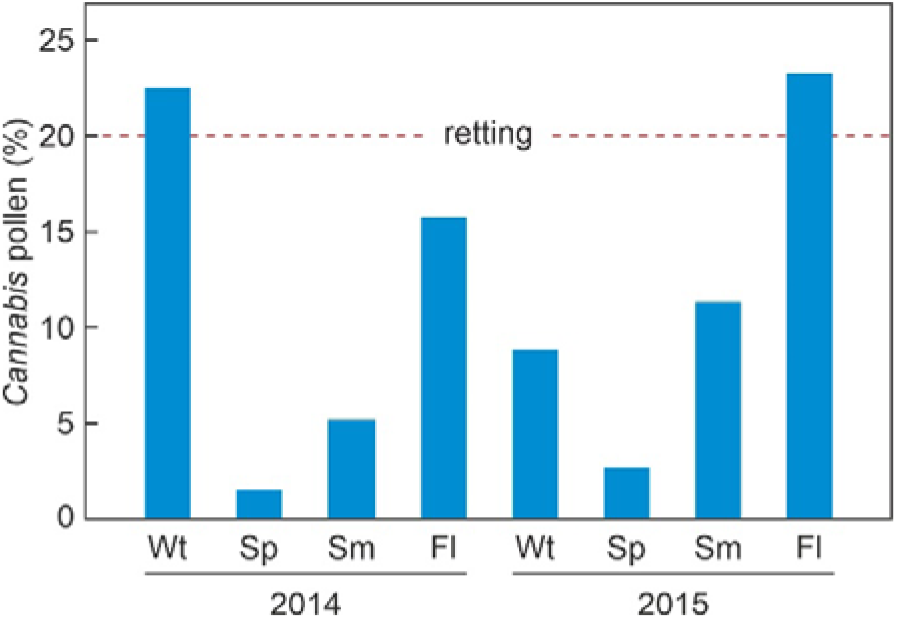
Seasonal distribution of sedimentary *Cannabis* pollen in Lake Montcortès during 2014 and 2015 (Rull et al., 2017). The hemp retting threshold (20%) is indicated by a red dashed line. Wt. winter; Sp, spring; Sm, summer; Fl, fall.

## 4. Discussion

As mentioned above, wild or cultivated *Cannabis* plants have not been reported in Montcortès local and regional vegetation surveys (Carreras et al., 2005-2006; Mercadé et al., 2013), and hemp retting has not been practiced in the lake since the late 19^th^ century, as indicated by historical records and supported by the absence of bRNA (Rull et al., 2022). Therefore, a reasonable explanation for the 20C event seems to be long-distance pollen dispersal mediated by wind (anemophily). However, the possibility of local or regional illegal crops cannot be disregarded (Munuera et al., 2002). In both cases, a conspicuous increase in these activities would be needed to explain the 20C. Therefore, pollen dispersal patterns and historical trends of *Cannabis* cultivation are critical points to be addressed.

The high productivity and dispersal power of anemophillous *Cannabis* pollen are well known (e.g., Small & Antle, 2003; Li et al., 2017). On the IP, several studies have documented relatively high amounts of airborne *Cannabis* pollen in areas far from known plant sources. For example, a northern African origin for the airborne *Cannabis* pollen found in some localities from the southeastern IP has been proposed by several authors after detailed aerobiological studies including the use of long-distance dispersal modeling tools. These localities include the cities of Murcia (Munuera et al., 2002; Aznar et al., 2022), Málaga (Cabezudo et al., 1997; Aboulaich et al., 2013) and Córdoba (Cariñanos et al., 2004) (Fig. 1), which are situated 150-300 km north of the African coasts. Pollen from northern Africa comes primarily from illegal crops (Aboulaich et al., 2013). The possibility of nearest sources, such as illegal local/regional crops, was suggested by Munuera et al. (2002). Recently, Aznar et al. (2022) found a significant increase in airborne *Cannabis* pollen between 2017 and 2020 in the region of Murcia (Fig. 1), which coincided with the predominance of local/regional wind patterns, and attributed this increase to the spread of illegal crops in the region rather than to long-distance dispersal from Africa. The occurrence of high *Cannabis* pollen percentages in modern sediments near crop fields of this plant has recently been documented (Boutahar et al., 2023). A typical feature of these crops is their high mobility and spatial turnover, which is a strategy to avoid detection and makes their identification difficult (Aboulaich et al., 2013). These studies come from the southernmost part of the IP, and no comparable information exists for other regions. However, it is expected that the dispersal of *Cannabis* pollen follows similar spatial patterns. Similar surveys should be conducted in the Montcortès region to evaluate the potential contribution of long-distance vs. local/regional pollen sources. These studies could also contribute to the detection of illegal cannabis crops in the area.

Regarding production, hemp cultivation almost disappeared from Spain in the early 1970s, but a renewed interest in this crop began in 1972. Until the 1960s, hemp was cultivated mainly on irrigated semiarid areas from the eastern IP (Alicante) to provide thread, ropes and fabric (Fig. 1). In 1972, however, the paper pulp industry started using hemp as a raw material instead of worn textiles, and in the early 1980s, the main production center shifted from the east to the northeast (Catalonia) in lowland areas (400-900 m elevation) of higher precipitation (600-700 mm per year), thus avoiding the need for continuous irrigation (Gorchs & Lloveras, 2003; Gorchs et al., 2017). This geographical shift coincided with a considerable production increase (<1000 tons in 1980 to >8000 tons in 1998) favored by the onset of EU subsidies to hemp cultivation, which started in 1985 (40 € ha^-1^) and significantly increased (∼780 € ha^-1^) in 1996-97 (Fig. 4). This production increase was centered on the southern Pyrenees, where Lake Montcortès (slightly above 1000 m elevation) lies, under wet climates, as a spring crop in rotation with winter wheat. This rotation significantly increased wheat yields, which has been attributed to the improvement of soil structure and the contribution of hemp to control pests, diseases and particularly weeds, as its rapid early growth outcompetes most weed species, thus reducing the weed burden for the ensuing winter crops (Gorchs et al., 2017).

**Figure 4.**
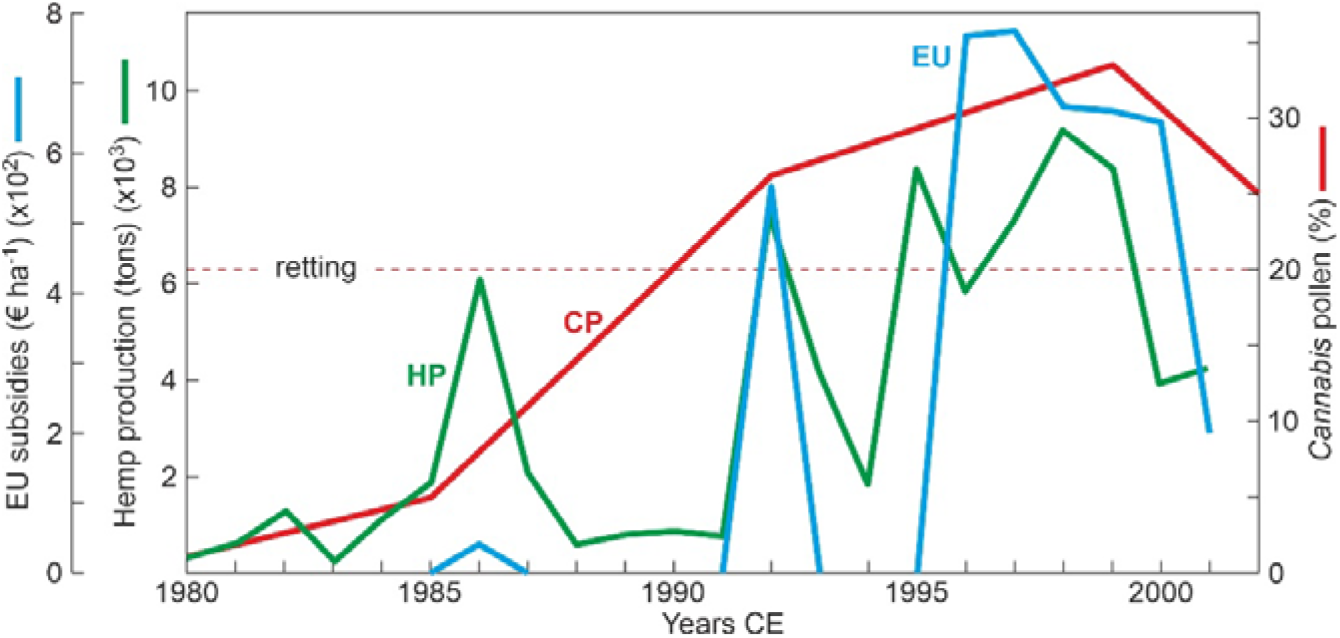
Hemp production in NE Spain (HP) and European Union subsidies to hemp cultivation (EU) (raw data from Gorchs & Lloberas, 2003) compared with the *Cannabis* pollen percentage curve from Lake Montcortès sediments (CP) (Fig. 2).

The 20C event, as recorded in Lake Montcortès was coeval with this 1980s hemp production acceleration, which strongly suggests a causal relationship (Fig. 4). Due to its high dispersion power, hemp pollen could have easily reached Lake Montcortès from adjacent regional pre-Pyrenean hemp crops, which could be confirmed by modeling the *Cannabis* pollen dispersal around the lake area. Hemp cultivation declined again shortly after the 1990s and was abandoned by 2006, due to the relocation of the hemp straw processing industry (Gorchs et al., 2017). This decline was also paralleled by pollen trends (Fig. 2), thus reinforcing the potential causal relationship. However, *Cannabis* pollen remained close to the retting threshold until at least 2015 (Figs. 2 and 3), which deserves an additional explanation. Additional hemp pollen sources from illegal crops cannot be disregarded, although the available evidence is less robust. In recent decades, a relevant rise in large-scale *Cannabis* plantations has been recorded in the IP, especially along the Mediterranean coasts, in a trend to increasingly replace hashish imports from northern Africa (Alvarez et al., 2016; Belackova et al., 2016). The potential manifestation of this peninsula-wide rising trend in the Montcortès region remains to be documented, and is worth to be addressed in future studies.

The geographical extent of the 20C event remains unknown, as it has not been recorded in other IP localities. This lack of evidence could suggest that this event could be a local/regional phenomenon, but methodological constraints should not be dismissed. On the one hand, only 23 of the 56 records included in the peninsular-wide CHIP database (Rull et al., 2023) contain the 20^th^ century, and of them, *Cannabis* pollen occurs in only four (Fig. 1), excluding Lake Montcortès. Of these four sites, *Cannabis* pollen is abundant (maximum of 25%) in only one (ET), whereas in the others this pollen has maxima of only 5% (CE) or shows single scattered occurrences (AD, EN). On the other hand, the resolution of these records, usually centennial to millennial, is not enough to resolve eventual changes that occurred within the 20^th^ century. For example, in ET, which is the most similar to Montcortès regarding pollen abundance, the record ends in 1991, and only a couple of samples are available for the whole 20^th^ century (Riera et al., 2004). Therefore, the lack of high-resolution records including the 20^th^ century is a major handicap to assessing the spatial patterns of the 20C across the IP.

Another interesting observation is that the major vegetation shift documented above and the 20C occurred at or shortly after the onset of the “Anthropocene”, as currently defined by the Anthropocene Working Group (AWG) (Waters et al., 2023). Indeed, the AWG proposes that this boundary should be situated in the mid-20^th^ century, coinciding with the Great Acceleration, when the Earth System underwent a global transformation overcoming the range of variability of the Holocene (Steffen et al., 2015). Our results, together with the very high resolution (seasonal) of the Montcortès record, suggest that the sediments of this lake could be a suitable candidate to identify and precisely date an eventual “Anthropocene” GSSP (Global Stratotype Section and Point) (Waters et al., 2018). The proxies identified by the AWG as the best suited stratigraphic markers (radioisotopes, fly ash, stable carbon and nitrogen isotopes) have not yet been analyzed in the Montcortès record, but efforts in this direction are worth doing, especially after the detection of significant Hg and Pb increases during the Industrial Revolution and the Great Acceleration in these sediments (Corella et al., 2017).

## 5. Conclusions and future prospects

A distinct and characteristic increase in *Cannabis* pollen has been recorded in the unique varved record of Lake Montcortès during the 1980s, which has been related to the significant increase in hemp production documented by the same time in the southern Pyrenees, fostered by the incoming EU subsidies to this crop. Additional sources for this pollen type, notably illegal cannabis plantations, cannot be disregarded but sound evidence is still lacking. The geographical extent of this palynological event, called here 20C (20^th^ century *Cannabis* increase), remains unknown due to the lack of similar continuous and absolutely dated high-resolution records for the 20^th^ century. A number of future studies are proposed to complement this preliminary assessment, including (i) increasing the resolution of the available pollen record from subdecadal (∼5 years) to annual, (ii) modeling the *Cannabis* pollen dispersion in the region around Montcortès, (iii) surveying the occurrence of illegal plantations as potential local sources for *Cannabis* pollen, (iv) increasing the number and geographical extent of 20^th^ century high-resolution records similar to Montcortès, and (v) evaluating the Lake Montcortès record as a potential candidate for the identification of the current AWG “Anthropocene” prospect.

## Acknowledgments

This work has been funded by the Ministry of Economy and Competitiveness, projects CGL 2012-3665 and CGL 2017-85682-R.

